# A cell fate switch in the *C. elegans* seam cell lineage occurs through modulation of the Wnt asymmetry pathway in response to temperature increase

**DOI:** 10.1101/849174

**Authors:** Mark Hintze, Sneha L. Koneru, Sophie P.R. Gilbert, Dimitris Katsanos, Michalis Barkoulas

**Affiliations:** Department of Life Sciences, Imperial College, London SW7 2AZ, United Kingdom

**Keywords:** seam cells, epidermis, temperature, Wnt pathway, cryptic genetic variation, robustness

## Abstract

Populations often display consistent developmental phenotypes across individuals despite the inevitable biological stochasticity. Nevertheless, developmental robustness has limits and systems can fail upon change in the environment or the genetic background. We use here the seam cells, a population of epidermal stem cells in *Caenorhabditis elegans*, to study the influence of temperature change and genetic variation on cell fate. Seam cell development has mostly been studied so far in the lab reference strain (N2), grown at 20° temperature. We demonstrate that an increase in culture temperature to 25°, introduces variability in the wild-type seam cell lineage with a proportion of animals showing an increase in seam cell number. We map this increase to lineage-specific symmetrisation events of normally asymmetric cell divisions at the final larval stage, leading to the retention of seam cell fate in both daughter cells. Using genetics and single molecule imaging, we demonstrate that this symmetrisation occurs via changes in the Wnt asymmetry pathway, leading to aberrant Wnt target activation in anterior cell daughters. We find that intrinsic differences in the Wnt asymmetry pathway already exist between seam cells at 20° and this may sensitise cells towards a cell fate switch at increased temperature. Finally, we demonstrate that wild isolates of *C. elegans* display variation in seam cell sensitivity to increased culture temperature, although seam cell numbers are comparable when raised at 20°. Our results highlight how temperature can modulate cell fate decisions in an invertebrate model of stem cell patterning.

## Introduction

During development, organisms must withstand environmental and genetic perturbations to produce consistent phenotypes (Felix and Barkoulas 2012). These phenotypes are often a product of complex developmental events that require a tight balance between cell division and cell differentiation (Soufi and Dalton 2016). A key example is stem cell divisions, consisting of highly controlled asymmetric and symmetric patterns, which are vital for generating cell diversity, as well as maintaining cell numbers in tissues and organs (Morrison and Kimble 2006; Knoblich 2008). Developmental robustness has inherent limits and certain perturbations can push a system outside its buffering zone (Braendle and Felix 2008; Barkoulas *et al*. 2013). In these cases, it is also important to understand how systems fail by investigating how perturbations precisely modulate developmental processes. Here we address the question of how changes in environmental temperature can affect cell fate outcomes using the nematode *C. elegans* as a model system. While it is well known that increasing or decreasing environmental temperature can change the development speed in *C. elegans*, the effect of temperature on specific cell division and fate acquisition events is less well understood. The *C. elegans* adult hermaphrodite consists of 959 somatic cells with their complete and stereotypical lineage characterised (Sulston and Horvitz 1977); this, alongside the isogenic nature of *C. elegans* populations, make it an attractive model to study environmental effects on development.

We focus here on the seam cells, which are a population of epidermal cells that are found along the two lateral sides of the animal body. Seam cell development has been used as a system to study mechanisms of stem cell patterning in an invertebrate model (Joshi *et al*. 2010). This is because seam cells show stem cell behaviour during larval development as they go through reiterative asymmetric divisions, where usually the posterior daughter retains the seam cell fate, while the anterior daughter differentiates to a neuroblast or acquires hyp7 fate and joins the syncytial epidermis (also known as hypodermis) (Figure 1A) (Chisholm and Hsiao 2012). *C. elegans* hatch with 10 seam cells on each lateral side and during the second larval (L2) stage a symmetric division increases the total seam cell number from 10 cells to 16 (Figure 1A). The exact pattern of seam cell divisions differs between each lineage in the head (H0-H2 cells), mid-body (V1-V6) and tail (T) region and over developmental time (Figure 1A). The balance between seam cell proliferation and differentiation is controlled through transcription factor activity (Koh and Rothman 2001; Cassata 2005; Nimmo *et al*. 2005; Kagoshima *et al*. 2007; Huang *et al*. 2009; Brabin *et al*. 2011; Gorrepati *et al*. 2013; Hughes *et al*. 2013) and the Wnt/β-catenin asymmetry pathway (Mizumoto and Sawa 2007b; Sawa and Korswagen 2013; Gorrepati *et al*. 2015), which is an adaptation of the conserved canonical Wnt signalling pathway in the context of an asymmetric division. In this case, selective activation of Wnt-dependent transcription in one of the two seam cell daughters relies on asymmetric localisation of Wnt components in mother cells that are polarised before division (Takeshita and Sawa 2005; Goldstein *et al*. 2006; Mizumoto and Sawa 2007a; Gleason and Eisenmann 2010; Baldwin *et al*. 2016).

**Figure 1:**
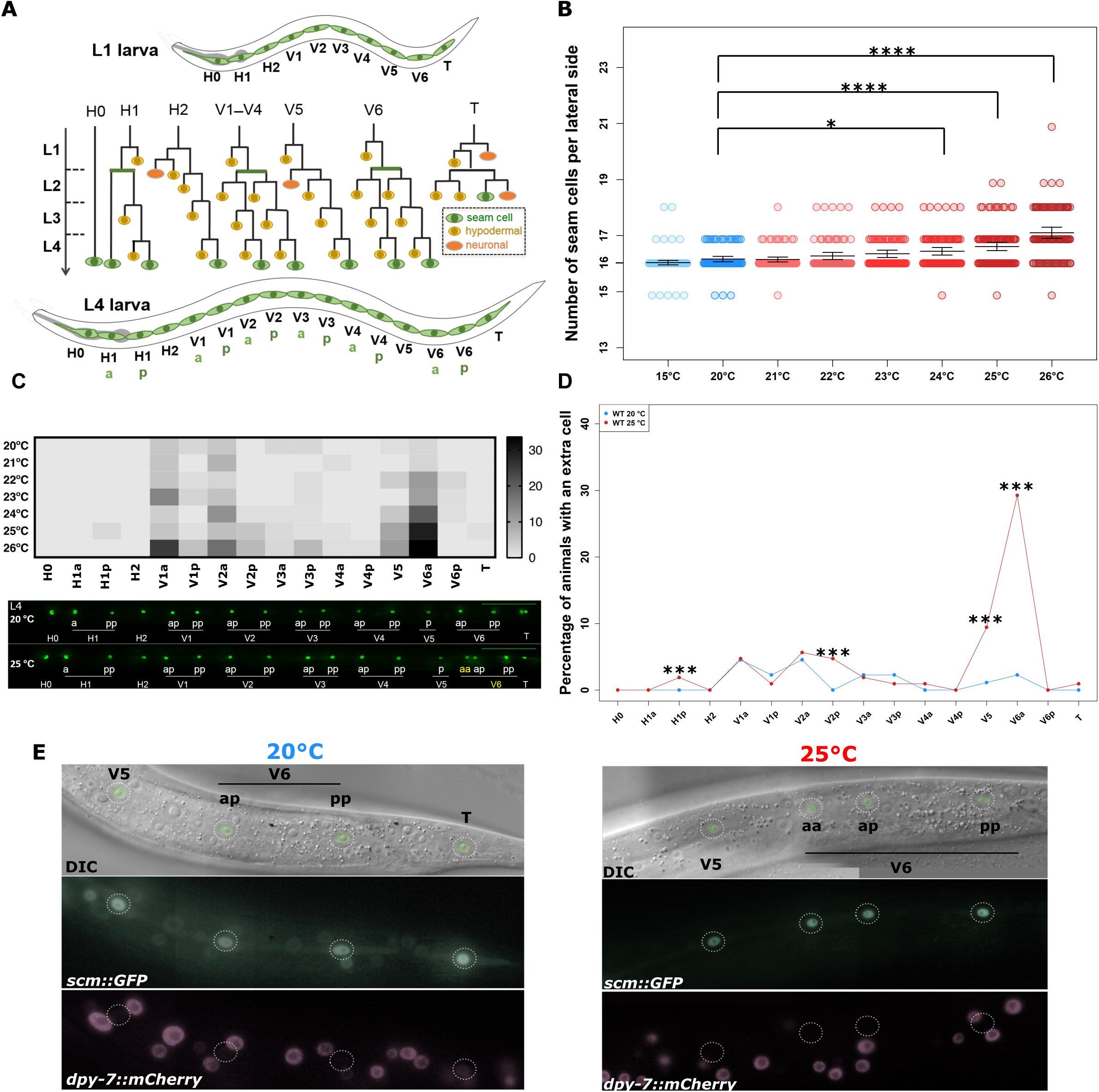
Lineage-specific seam cell duplications occur in higher growth temperature. **(A)** Schematic showing WT seam cell division patterns across larval stages (green = seam cell, yellow = hypodermal cell, orange =neuroblast). (**B)** Counts of seam cell nuclei in JR667 (*wIs51*[*scm::GFP*] used as WT reference) when animals were grown at temperatures from 15° (blue dots) to 26° (red dots) (* corresponds to *P*<0.05, **** *P*<0.0001, with one-way ANOVA followed by Dunnett’s multiple comparison test, n ≥ 79). Error bars represent 95% confidence interval of the mean. **(C)** Heat map displaying occurrence of errors along the seam axis as temperature increases (upper panel). Lower panel shows representative images of animals grown at 20° and 25°, yellow labels indicate extra seam cells in the V6a lineage at 25°. Scale bar is 100 µm. **(D)** Quantification of percentage of seam cell duplications per individual seam cell. Note that the highest proportion of duplications was seen in V6a at 25° (\*\*\**P*<0.001 with a binomial test, n = 88 and 106, 20° and 25° respectively). **(E)** In instances of extra nuclei in the V6a lineage, anterior V6a daughter cells (V6aa) do not express the hypodermal marker *dpy-7p::mCherry* and retain *scm::GFP* expression.

Despite progress made over the last years in identifying key factors contributing to epidermal development, most studies have been conducted using a single *C. elegans* isolate grown in a single environment - that is the lab reference strain N2 grown at 20°. It remains therefore unknown whether changes in the growth environment or the genetic background would have an impact on seam cell patterning. In this study, we start to address this question by investigating the effect of different growth temperatures, as well as genetic backgrounds, on seam cell development. We demonstrate that as culture temperature is increased within physiological limits (e.g. 25°), populations become more variable and start producing one extra seam cell on average. We show that this increase in seam cell number occurs via a cell fate switch that is observed reproducibly in specific cell lineages. This cell fate switch is dependent upon the Hox gene *mab-5* and the beta-catenin gene *bar-1,* both previously unknown to play a role in the hermaphrodite seam cell lineage at 20°. We show that at high temperatures, an impaired Wnt asymmetry pathway leads to ectopic Wnt pathway activation in anterior daughters of specific seam cells that may already be sensitised regardless of the growth temperature. Finally, we study here seam cell development for the first time outside N2 and find that wild isolates of *C. elegans* show a conserved seam cell number at 20°. Nevertheless, by raising animals at 25°, we reveal cryptic genetic variation between isolates and show that the sensitivity of the seam cell lineage to higher temperature evolves within *C. elegans,* with certain isolates showing an enhanced or suppressed response in comparison to N2. Together, these findings expand our knowledge of developmental system behaviour upon environmental and genetic perturbation.

## Materials and Methods

### Nematode culture and genetics

Strains used in this study were maintained and handled according to standard protocols (Brenner 1974). The strain JR667 containing a *scm::GFP* transgene (*wIs51*) in the N2 background is used as a reference stain and was maintained with OP50 as food source on standard NGM plates. The *scm::GFP* marker was introgressed from JR667 into wild isolates JU775, JU2519 and CB4856 together with a *dat-1::GFP* marker using a two-step cross repeated five times to produce ten times backcrossed strains. In the first step, hermaphrodites carrying the *vtIs1*[*dat-1p::GFP*] *+ wIs51*[*scm::gfp*] transgenes linked on chromosome V were mated to wild-isolate males. In the second step, F1 males from the previous cross were crossed to wild-isolate hermaphrodites. F1 hermaphrodites carrying both *wIs51* and *vtIs1* were allowed to self and homozygous progeny for the marker were considered backcrossed twice. The linkage between the two transgenes (*vtIs1* and *wIs51*) was broken at the last step when F1 hermaphrodites, from a cross between wild-isolate hermaphrodites carrying these transgenes and wild isolate males, were allowed to self. Recombinant progeny carrying only *wIs51* were picked and maintained. *scm::GFP* was introgressed into wild isolates (JU2007 and XZ1516) by crossing wild isolate males to JR667 hermaphrodites. F1 males carrying the transgene were crossed to wild isolate hermaphrodites. This last step was repeated nine times to produce ten times backcrossed wild isolates. A complete list of strains used in this study is presented in Table S1.

### Microscopy and phenotypic characterisation

Standard seam cell scorings were performed by mounting animals on 2% agar pads containing 100 µM sodium azide (NaN_3_). These were covered with a coverslip and viewed under a compound microscope (AxioScope A1; Zeiss). Seam cell numbers were scored at the late L4 or early adult stage focusing on one lateral side per animal.

To perform temperature treatments, synchronised animals were prepared by bleaching adults and placing eggs on standard NGM plates at different temperatures. To quantify seam cells, animals were collected between 44 and 48 hours for L4s grown at 20° and between 36 and 40 hours for L4s at 25°. Seam cell duplications were scored based on the stereotypical position of seam cells in relation to the vulva and in relation to each other. More specifically, eight seam cells (H0, H1a, H1p, H2, V1a, V1p, V2a, V2p) are anterior to the vulva, two are adjacent to the vulva (V3a and V3p) and six seam cells are found posterior to the vulva (V4a, V4p, V5, V6a, V6p, T). In cases of increased seam cell number, the extra seam cell was assigned, when possible, to the nearest seam cell neighbour at the L4 stage, taking also into account the position of the corresponding seam cells on the opposite lateral side. Throughout this manuscript, we refer to the anterior and posterior branch of a seam cell lineage at a given developmental stage as a and p and their daughters as aa/ap and pa/pp. For example, the V6a lineage at L4 includes V6pappa (simplified here as V6aa) and V6pappp (V6ap).

POP-1 levels were characterised using a strain carrying the transgene *qIs74*[*sys-1p:: GFP::POP-1*]. Images of seam cells were analysed using ImageJ. The following formula was used to calculate the corrected total cell fluorescence (CTCF) in cell nuclei: Integrated Density − (Area of selected cell × Mean fluorescence of background readings), with three background readings around the animal taken for each cell pair. Cells with an anterior/posterior intensity ratio above 1.1 or below 0.9 were classified as anterior > posterior or anterior < posterior respectively, while cell pairs that had a ratio between 0.9 and 1.1 were classified as equal in fluorescence intensity.

### RNAi by feeding

Animals were fed with dsRNA expressing bacteria as a food source. HT115 bacteria containing RNAi clones or an empty-vector control were grown overnight and seeded directly on NGM plates that contained 1 μM IPTG, 25 μg/ml ampicillin and 6.25 μg/ml tetracycline. All RNAi clones used in this study were sequence-validated and come from the Ahringer RNAi library (Source Bioscience).

### Cloning

A *dpy-7p::mCherry::H2B:unc-54* cassette was assembled in vector pCFJ906 using standard three fragment Gateway cloning (Invitrogen). A recovered mimiMos insertion was crossed to JR667 to generate strain MBA227. To construct a *pseam::GFP::CAAX::unc-54* transgene, the GFP sequence was amplified using pPD93.65 as template and fused to the following sequence containing the CAAX motif using nested PCR 5’AAGGACGGAAAGAAGAAGAAGAAGAAGTCCAAGACCAAGTGCGTCATCATG3’. The GFP::CAAX fragment was then subcloned into pIR5 (Katsanos *et al*. 2017) via Gibson assembly. A stable integrant was obtained via transgenesis and gamma irradiation. The resulting line was backcrossed ten times to N2 before crossing to JR667 to generate strain MBA237.

### Single molecule fluorescence *in situ* hybridisation

Populations of nematodes were synchronised by bleaching and subsequently fixed in 4% formaldehyde at appropriate stages for the experiment (17 hours to image L1s and 40 and 44 hours to capture the L4 division at 20° and 25° respectively). smFISH was performed as previously described (Katsanos *et al*. 2017) using a pool of 27 – 48 oligos fluorescently labelled with Cy5 (Biomers). Imaging was performed using a motorized epifluorescence Ti-eclipse microscope (Nikon) and a DU-934 CCD-17291 camera (Andor Technology, United Kingdom) acquiring 0.8 μm step z-stacks. Image analysis and spot quantification were performed on raw data using a MATLAB (MathWorks, Natick, MA) routine as previously described (Barkoulas *et al*. 2013). For all images presented in this study, the probe signal channel was inverted for clarity (black spots correspond to mRNAs) and merged to the seam cell channel (GFP) using ImageJ (NIH, Rockville, MD).

### Data Analysis and availability

Data were analysed and presented with the R programming environment or GraphPad Prism 7. Two-sample t-tests were performed for differences in mean seam cell number. Binomial tests were performed to test for differences in the proportion of seam cell symmetrisation events between strains and/or treatments. All statistics were carried out in R 3.2.0. All reagents and strains are available upon request. Nematode strains are listed in Table S1. Table S2 contains oligo sequences used as smFISH probes in this study. Table S3 contains raw counts from smFISH probes for figures 2, 3, 4 and S3.

**Figure 2:**
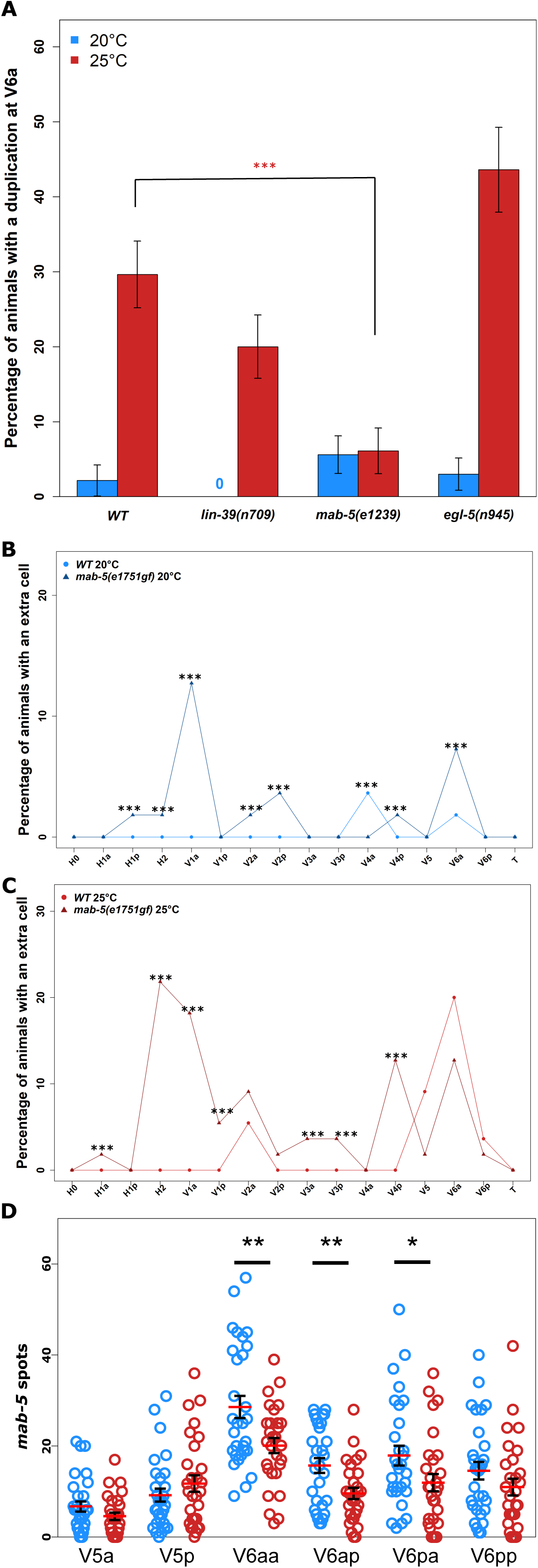
The Hox gene *mab-5* is necessary and sufficient for seam cell duplications. **(A)** The loss-of-function allele *mab-5(e1239)* suppresses V6a duplications when compared to WT (N2) at 25° (*P*<0.001 with a binomial test), while mutations in *lin-39(n709)* or *egl-5(n945)* do not display a similar effect, n≥80. Error bars indicate standard error of the proportion. **(B-C)** *mab-5* gain-of-function mutants (*mab-5e1751gf,* triangle points) show significant increase in duplications across seam cells both at 20° and 25° compared to WT (circle points) animals (*** correspond to *P*<0.001 with a binomial test, n≥40). **(D)** *mab-5* mRNA levels in V5, V6a and V6p daughter cells measured by smFISH in animals grown at 25° (red circles, n= 28) compared to 20° animals (blue circles, n=29). Levels of *mab-5* are significantly decreased in V6aa, V6ap and V6pa (* *P*<0.05, ** *P*<0.01 with a two-sample t-test). Error bars indicate standard error of the mean.

**Figure 3:**
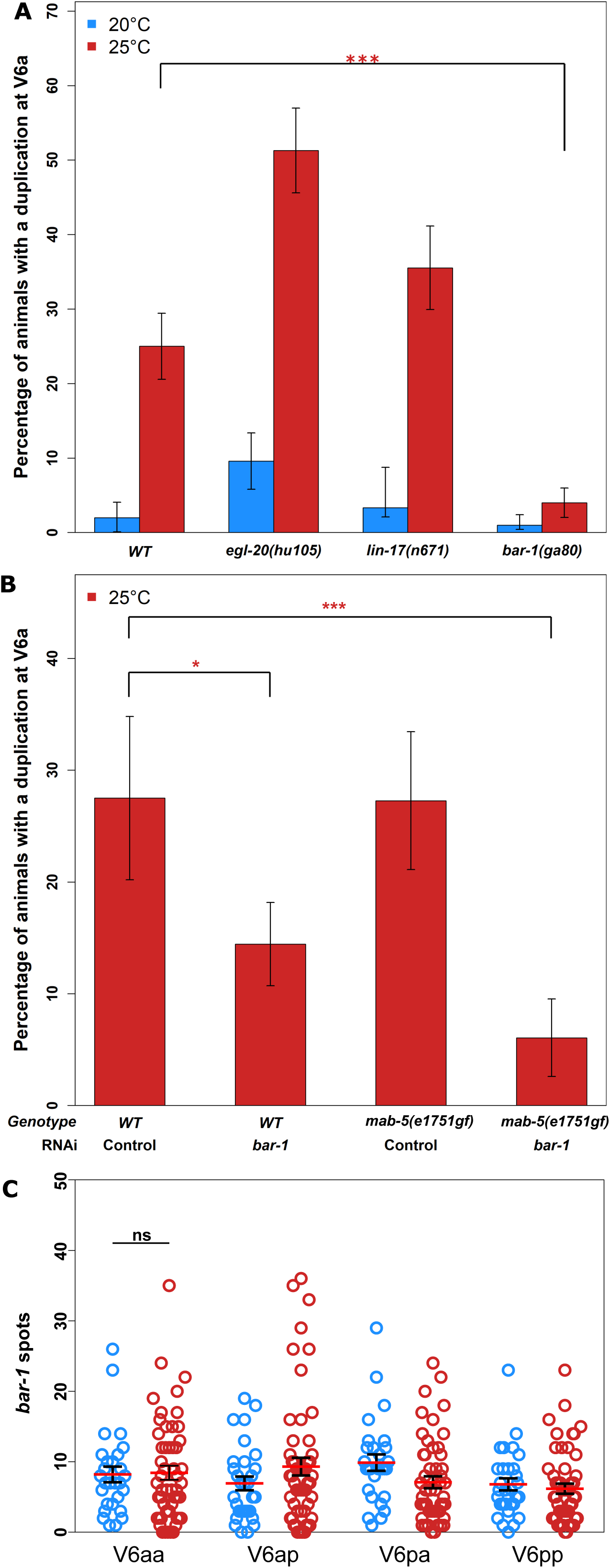
The beta-catenin *bar-1* is necessary for V6a duplications. **(A)** V6a duplications were significantly suppressed in *bar-1(ga80)* mutants at 25° compared to wild-type animals grown at 20° (*** *P*<0.001, binomial test, n≥90). No suppression was seen in *egl-20(hu105)* and *lin-17(n671)* mutants, n≥30. **(B)** The proportion of animals displaying V6a seam cell duplications in WT and *mab-5(e1751gf)* animals was significantly decreased when grown at 25° on *bar-1* RNAi (* *P*<0.05, *** *P* <0.001, n ≥40). **(C)** mRNA levels of *bar-1*, measured by smFISH, are not significantly different in animals grown at 20° and 25° in V6a and V6p daughter cells (n = ≥ 25). Error bars indicate standard error of the proportion (A, B) or mean (C).

**Figure 4:**
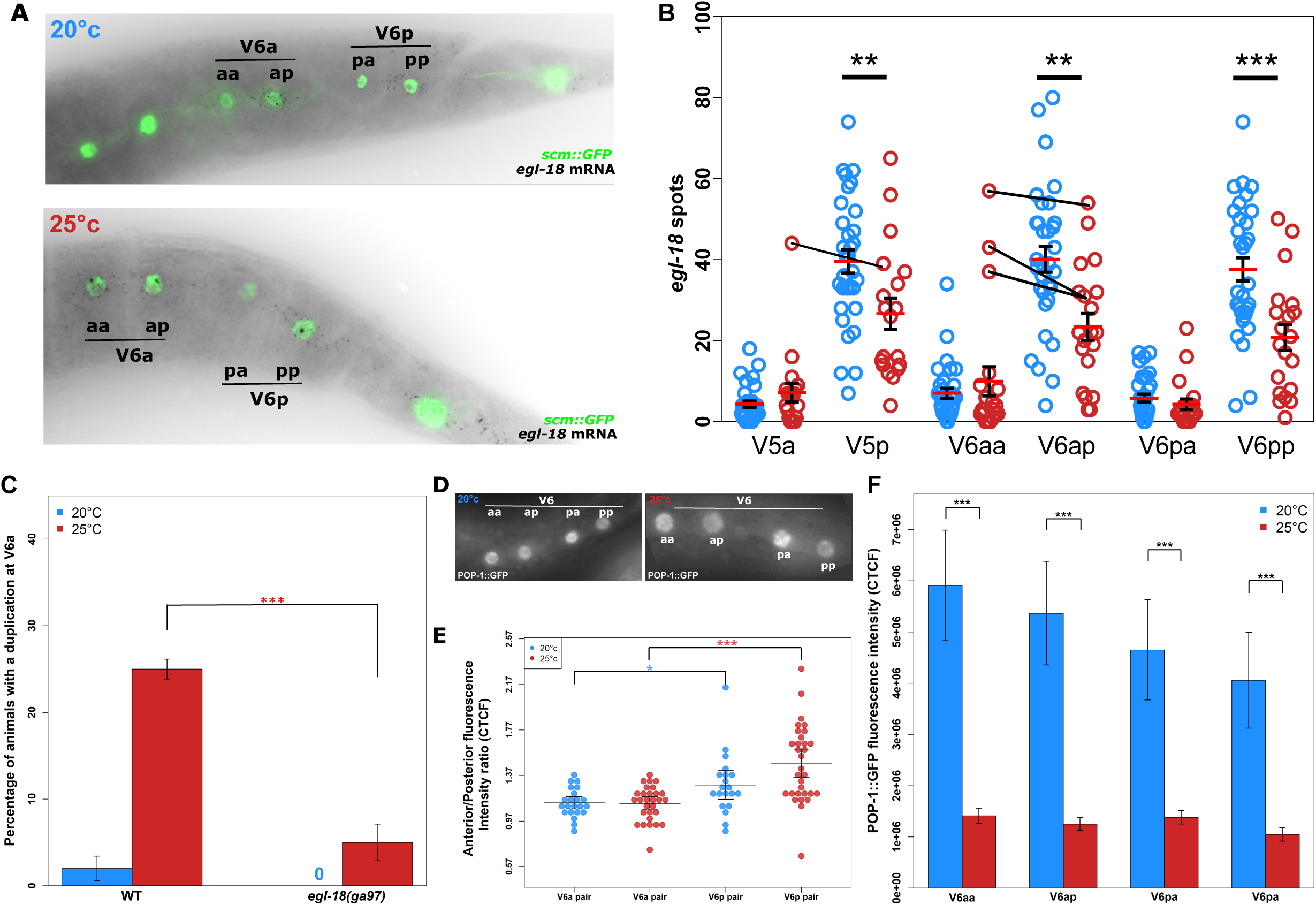
Wnt pathway asymmetry is impaired in V6a daughters. **(A)** Representative images of *egl-18* smFISH (black dots correspond to mRNAs) in V6a and V6p daughter cells at 20° and 25°. Seam cells appear green due to expression of the *scm::GFP* marker. **(B)** Quantification of *egl-18* mRNA levels in V5, V6a and V6p daughters at 20° and 25°. Expression was significantly lower in posterior cells (V5p, V6ap, and V6pp) at 25° (red circles) compared to the same seam cells grown at 20° (blue circles, ** *P <*0.01, *** *P <*0.001 with a two-sample t-test, n>20 per cell). At low frequency, animals grown at 25° showed extreme expression values in anterior V6aa cells and V5a, which were higher than their posterior counterparts (these pairs are indicated by black lines). **(C)** Seam cell duplications at V6a are suppressed in the *egl-18(ga97)* mutant at 25°, (*** *P*<0.001 with a binomial test, n = 120). (**D)** Representative images of nuclear POP-1::GFP expression at 20° and 25°. **(E)** The ratio of nuclear POP-1::GFP expression between V6a and V6p daughter cells at 20° and 25°, n≥20 (\**P*<0.05, *** *P*<0.001, two-sample t-test, error bars indicate 95% confidence interval of the mean. **(F)** Average POP-1::GFP intensity in V6a and V6p daughter cells at 20° and 25° (error bars represent standard error of the mean, * *P*<0.05, ** *P*<0.001, with a two-sample t-test).

## Results

### Increase in growth temperature leads to extra seam cells in specific lineages

Seam cell development has been mostly studied so far at the standard growth temperature of *C. elegans* in the lab, which is 20°. We therefore decided to investigate whether varying the growth temperature could affect seam cell development. To this end, we cultured *C. elegans* at a range of temperatures ranging from 15° to 26° and scored seam cell number based on the expression of an *scm::GFP* marker (*wIs51*). We placed eggs to hatch at different temperatures and scored terminal seam cell number at the end of the fourth larval stage (L4). At this stage, all somatic divisions are completed, and so terminal seam cell number acts as a potential read-out for seam cell defects that have accumulated over post-embryonic development.

As expected, populations grown at 20° showed an average seam cell number of approximately 16 cells per lateral side and low phenotypic variance because of the rare occurrence of animals displaying 15 or 17 seam cells (Figure 1B). While we found no statistically significant difference in seam cell number when the culture temperature was decreased to 15°, we were surprised to see that populations grown at 23° and above showed a mild, yet statistically significant, increase in terminal seam cell number (Figure 1B, *P*<0.05, two-sample t-test). This increase, due to the frequent occurrence of animals displaying 17 seam cells instead of the wild-type 16, was most frequent at 25° and 26°. We decided to use 25° for all subsequent experiments, which is considered to be a viable physiological temperature for *C. elegans* growth, and is commonly used as an alternative temperature to 20° for example in order to accelerate development or study temperature-sensitive mutants.

We first sought to investigate the developmental basis underlying the increase in seam cell number at 25°. While scoring seam cell number at different temperatures, we observed that a frequent error at 25° was the tight clustering of two posterior seam cell nuclei, a phenotype that we refer to here as “seam cell duplication” for simplicity. We mapped the frequency of extra seam cells along the axis of the larva and assigned them to seam cell lineages based on their position relative to the position of their closest seam cell neighbour. This highlighted a significant hotspot for seam cell duplications at the anterior V6 lineage (simply here symbolised as V6a but formally V6pappp/V6pappa), with around 30% of animals in the population showing this phenotype when grown at 25° in contrast to only 2% at 20° (Figure 1C and D, *P*<0.001, binomial test). Seam cell duplications were not exclusive to the V6a lineage, but were also observed, albeit with lower frequency, in the V5, V1a and V2 lineages (Figure 1C and D). This finding indicates that the various seam cell lineages display different sensitivities to temperature increase.

To understand when these duplications occur during post-embryonic development, we transferred animals at different developmental stages from culture at 20° to 25°. We found that transferring animals at any stage before L2 resulted in a similar increase in seam cell number, suggesting that seam cell lineage errors occur after the L2 developmental stage (Figure S1A). By scoring the frequency of extra V6a cells at the later larval stages in animals raised entirely at 25°, we found that seam cell duplications occurred during the L4 division (Figure S1B). This observation is consistent with the close clustering of pairs of nuclei observed at the end of the L4 stage, indicative of a recent developmental event.

The seam cell duplications observed may be a consequence of defects in cell division or cell differentiation during L4. With regard to cell division defects, one possibility is that the extra nuclei are a result of a failure of cytokinesis at 25°. To address this possibility, we used a marker of seam cell membranes to look for multinucleate cells at L4, but we did not observe any defects in cytokinesis in instances of extra nuclei in the V6a lineage (Figure S1C). Although we have not formally ruled out the possibility of an ectopic seam cell division, we believe this is unlikely due to the number of animals we have observed throughout the L4 stage at 25°, during which we have never observed evidence of a cell division in addition to the wild-type L4 seam cell division pattern. We then explored whether, despite the seam cells dividing successfully, the anterior daughters fail to differentiate into hyp7, but instead retain the seam cell fate. Indeed, using a strain that carries both the seam cell and a hyp7 cell marker (*dpy-7p::mCherry*), we found that both cells of the duplicated V6a lineage expressed the seam cell marker alone, with neither cell expressing the hyp7 cell marker (Figure 1E). Finally, we reasoned that the additional seam cell may reflect a timing constraint for V6a daughters to differentiate before terminal seam cell fusion occurs, since seam cells terminally fuse at the end of the L4 stage and development is accelerated at 25°. We argue that this is unlikely to be the case because we found that V6a is not the last seam cell to divide at L4, despite displaying the highest sensitivity to temperature (Figure S1D). Furthermore, V6a duplications were still observed in an *aff-1* mutant background, which is impaired in terminal seam cell fusion (Figure S1E). Taken together, these results indicate that an increase in culture temperature induces seam cell duplications during the L4 division, due to conversion of asymmetric cell divisions to symmetric wherein both cells adopt the seam cell fate.

### The Hox gene *mab-5* is necessary for seam cell duplications at 25°

Ectopic seam cells at 25° were most frequently found in specific lineages, which highlights differences in sensitivity to temperature along the nematode body axis. We therefore decided to investigate whether factors involved in anterior-posterior patterning can influence the seam cell phenotype in response to temperature increase. Hox genes are known to be involved in the specification of the animal body axis and play a role in seam cell patterning as well (Salser and Kenyon 1996; Arata *et al*. 2006). A posterior body Hox gene, *mab-5,* has been reported to be required for V5 and V6 seam cell lineages in males to generate sensory rays, while loss of *mab-5* leads to a switch from ray formation to alae (Salser and Kenyon 1996). To investigate the role of Hox genes on V6a lineage defects at 25°, we compared terminal seam cell number and frequency of V6a duplication in three strains carrying individual loss-of-function mutations for the mid and posterior Hox genes *lin*-*39*, *egl-5* and *mab-5* (Austin and Kenyon 1994; Salser and Kenyon 1996; Maloof and Kenyon 1998) (Figure 2A and Figure S2A). We found that only loss of *mab-5* function significantly suppressed V6a seam cell duplications at 25° (Figure 2A, *P*<0.001, binomial test), indicating that *mab-5* is necessary for V6a duplication events. In addition, we found that *mab-5* is required for the maintenance of posterior seam cell fates in hermaphrodites, as *mab-5* loss-of-function animals lost posterior seam cells at low frequency (Figure S3A). This seam cell loss is unlikely to affect the comparison between temperatures because the frequency of V6a loss was similar between 20° and 25° (Figure S3A). To investigate whether *mab-5* is also sufficient for seam cell duplications, we quantified seam cell number in a *mab-5* gain-of-function mutant *mab-5(e1751gf*). In this background, *mab-5* expression is thought to expand from the posterior to the anterior end due to a *mab-5* genomic locus duplication (Salser and Kenyon 1992). We confirmed *mab-5* expansion at the mRNA level with single molecule FISH (smFISH) (Figure S3, B-E). Interestingly, we found that *mab-5(e1751gf*) mutants show a significant increase in ectopic seam cells, which was observed even at 20° and became more pronounced at 25° with multiple seam cell lineages showing seam cell duplications (Figure 2, B and C, *P*<0.001, binomial test).

Based on the *mab-5* gain-of-function seam cell phenotype, we then hypothesised that an increase in *mab-5* mRNA levels may underlie the seam cell duplications we observed at 25°. To address this hypothesis, we quantified *mab-5* expression by smFISH in posterior seam cells in wild-type animals at the L4 stage. We focused on V5 and V6a cell daughters, which are very sensitive to fate change in higher temperature, in comparison to V6p daughters which are less sensitive. Surprisingly, we found a significant decrease in mRNA expression at 25° compared to 20° (Figure 2D and S3F, *P<0.05*, two-sample t-test), although the expression of *mab-5* showed a peak in V6aa (formally V6pappa) both at 20° and 25° degrees, which may relate to the sensitivity of this lineage to fate change upon temperature increase. Taken together, we conclude that *mab-5* is necessary and sufficient for seam cell lineage fate changes during L4. However, seam cell duplications in response to higher temperature are unlikely to be driven at the level of an increase in *mab-5* mRNA expression.

### Seam cell duplications require the canonical Wnt pathway component BAR-1

Due to the role of Wnt signalling in controlling seam cell division patterns, we went on to investigate whether mutations in candidate Wnt components may suppress the V6a seam cell symmetrisation at 25°. For example, the Frizzled receptor LIN-17 and the posteriorly produced Wnt ligand EGL-20, have previously been shown to polarise seam cell divisions and interact with *mab-5* during Q neuroblast migration and (Mizumoto and Sawa 2007b; Middelkoop and Korswagen 2014). However, we found that mutations in either of these factors were unable to supress the V6a duplication (Figure 3A and Figure S2B). Instead, loss of the canonical Wnt pathway beta-catenin *bar-1* led to a significant decrease in the number of V6 seam cell duplications at 25° compared to wild-type (Figure 3A, *P*<0.001, binomial test). To validate this result, we knocked-down *bar-1* by RNAi and found that this treatment significantly reduced the number of V6a duplications observed at 25° in wild-type (Figure 3B, *P<*0.05, binomial test). This *bar-1* RNAi treatment also suppressed the V6a duplications in the *mab-5(gf)* background (Figure 3B, *P*<0.001, binomial test), indicating that *bar-1* is likely to be required for posterior *mab-5* expression or act in parallel to *mab-5* to facilitate seam cell symmetrisation at 25°.

The dependence of seam cell duplication on *bar-1* was surprising as this gene was not previously thought to play a major role in seam cell development studied at 20°. We therefore tested if *bar-1* expression could be detected in seam cells, and subsequently if *bar-1* mRNA levels were changed when animals were cultured at 25°. We were able to detect *bar-1* expression by smFISH in both anterior and posterior V6 seam cell lineages at the L4 stage (Figure 3C). However, we found no significant change in mean number of *bar-1* transcripts in V6aa cells that show the seam cell duplication phenotype at 25° versus 20°. These results indicate that *bar-1* is expressed in seam cells at the time when seam cell asymmetric division defects occur and suggest that these defects may occur through a temperature-driven activation of the Wnt pathway in anterior seam cell daughters of the V6 lineage.

### Impaired Wnt pathway asymmetry underlies seam cell fate changes

One of the main pathways involved in maintenance of seam cell fate and regulation of the asymmetric cell division is the Wnt pathway (Gleason and Eisenmann 2010; Sawa and Korswagen 2013). Following an asymmetric seam cell division, activation of the Wnt pathway is usually restricted to posterior cell daughters, which express key downstream genes such as the GATA transcription factor *egl-18* and maintain the seam cell fate(Gleason and Eisenmann 2010; Gorrepati *et al*. 2013). Ectopic activation of the Wnt pathway in anterior cell daughters has been shown to be sufficient for these cells to adopt the seam cell fate, in a similar manner to their posterior counterparts. This occurs for example upon *pop-1/tcf* down-regulation and leads to a dramatic increase in the average seam cell number (Gleason and Eisenmann 2010). To address whether defects in asymmetric seam cell division were associated with changes in Wnt pathway activity localisation, we quantified *egl-18* mRNA expression in animals grown at 20° and 25° during the L4 division. As expected, we found that at 20° the posterior cell daughters of V5, V6a and V6p following the L4 asymmetric division all expressed *egl-18* at a higher level than their anterior sister cells (Figure 4, A and B). Consistent with our findings of ectopic seam fate retention at 25°, we found that a subset of anterior cells showed at low frequency expression values near or beyond those anticipated for posterior daughter cells. V6a anterior daughters in particular showed a noticeable increase in extreme *egl-18* expression values compared to 20° (Figure 4B, see black lines connecting anterior to posterior cell pairs). This shift towards high *egl-18* mRNA values in a proportion of V6aa cells at 25° is likely to underpin the seam cell duplications observed. In addition, we observed that the posterior V daughter cells that are fated to remain seam cells at 25° showed significantly less *egl-18* expression than the same cells at 20° (Figure 4B). These results are indicative of a molecular shift in the L4 division at 25° from an asymmetric towards a symmetric mode.

To test whether *egl-18* plays a functional role in the symmetrisation of L4 seam cell divisions at 25°, we scored seam cell duplications in an *egl-18* loss-of-function mutant. We focused on the V6a lineage, which is largely unaffected in this mutant background at 20°, to assess seam cell fate symmetrisation frequency at 25° and found that loss of *egl-18* activity suppresses the duplication frequency (Figure 4C, *P*<0.001, binomial test). Taken together, these data support that seam cell duplications may occur due to ectopic activation of Wnt pathway in anterior seam cell daughters at 25°.

Nuclear levels of POP-1 are a good indicator of post-division asymmetry in seam cells, with high POP-1 levels (usually in anterior daughters) associated with a non-seam cell fate due to repression of Wnt targets and lower levels (in posterior daughters) associated with activation of Wnt targets and retention of seam cell fate (Gleason and Eisenmann 2010). To investigate whether the distribution of POP-1 is changed in seam cell daughters at 25°, we used a strain carrying POP-1::GFP and compared the levels of nuclear GFP expression between V6a and V6p seam cell daughters during the L4 division. We found that a third of V6a cell pairs (V6aa-V6ap) had equivalent POP-1::GFP levels both at 20 and 25°. This was in contrast to V6p cell pairs (V6pa-V6pp) which maintained significantly higher levels of POP-1 expression in anterior versus posterior daughters both at 20° and 25° (*P*-value<0.05, two-sample t-test) (Figure 4, D and E). We also observed that the overall level of POP-1::GFP was significantly decreased in all V6a and V6p cell pairs at 25° during the L4 division (Figure 4F). These results suggest that V6 cells may have intrinsic differences in Wnt pathway asymmetry, which make them more sensitive to temperature perturbations. When this is combined with a lowering in the overall amount of nuclear POP-1 in V6 lineage cells, this sensitivity may lead to a greater chance of anterior V6a daughter cells retaining a seam cell fate.

### The genetic background modifies the pattern and frequency of seam cell fate changes at 25**°**

Seam cell development has never been studied in any other *C. elegans* isolate except for the lab reference strain N2. Over the last few years, several divergent *C. elegans* strains have been isolated from various locations throughout the world, offering now the opportunity to study the effect of the genetic background on seam cell patterning and its robustness to various perturbations, including temperature increase.

We sought here to investigate whether seam cell number is robust to standing genetic variation and whether the observed seam cell duplications in response to higher temperature could also be observed in other natural isolates. To be able to visualise the seam cells, we first genetically introgressed the seam cell marker *scm::GFP* into five wild isolates by repeated backcrossing. We included strains which are known to be significantly divergent from N2, such as the commonly used Hawaiian isolate CB4856 (Andersen *et al*. 2012). We found that all five isolates we tested had an average of 16 seam cells per lateral side at 20°, which is the same as N2 (Figure 5A and Figure S4A). However, they responded differently to temperature increase (Figure 5A and Figure S4A). In particular, isolate XZ1516 was much more sensitive and showed higher frequency of duplications in various seam cell lineages when cultured at 25° (Figure 5B and Figure S4, B-D). Isolate JU2519 showed duplications in lineages (V3) that do not show cell fate defects in N2. Interestingly, V6a remained the most sensitive cell lineage to temperature increase in all isolates that were sensitive to temperature. On the other end of the spectrum, seam cells in one isolate (CB4856) were robust to temperature increase, with the frequency of seam cell lineage defects being the same between the two growth temperatures of 20° and 25° (Figure 5, A and B). Together, these results indicate that variation in the genetic background can both enhance and suppress the seam cell fate changes observed upon temperature increase.

**Figure 5:**
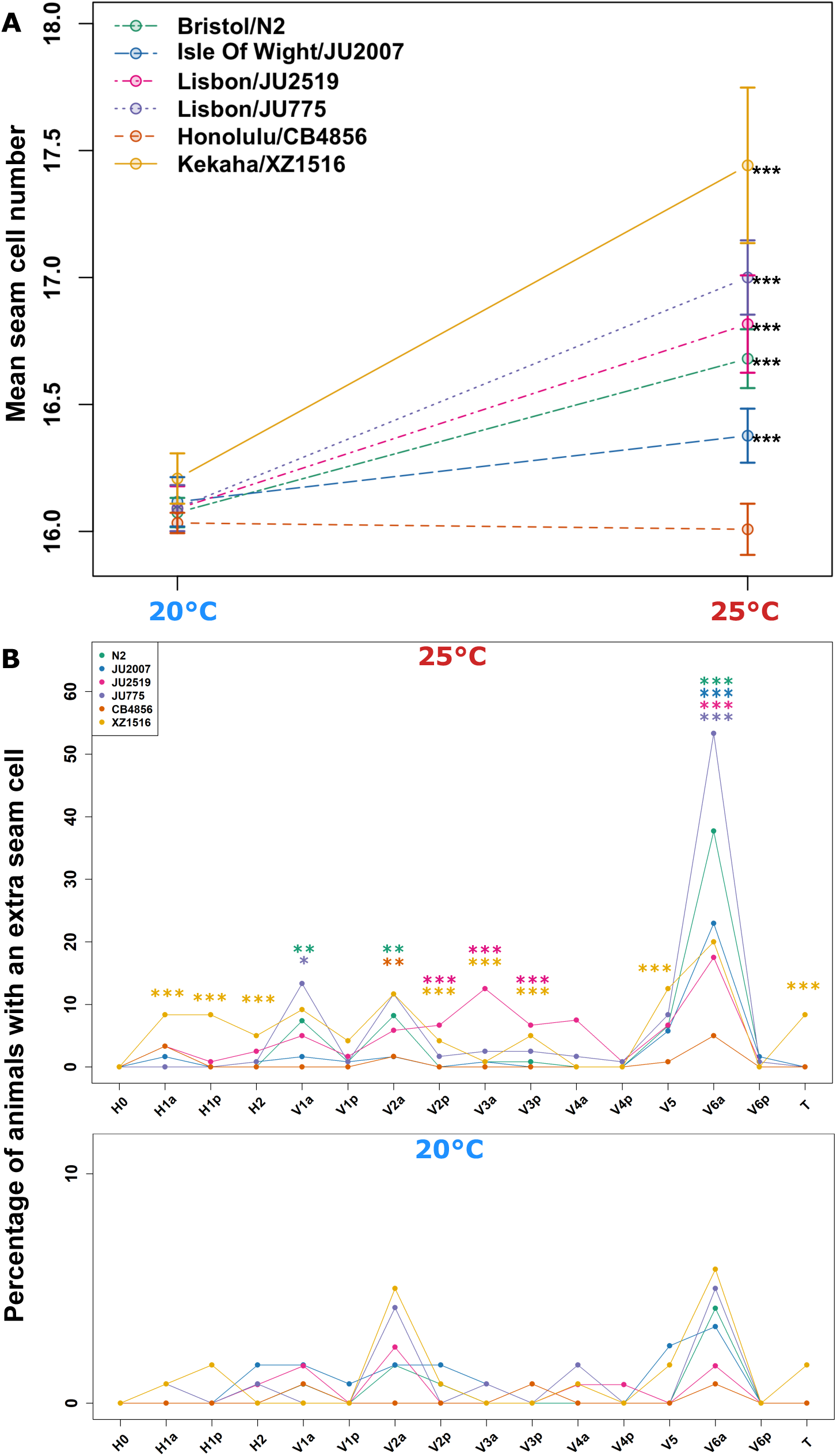
Evolution of the frequency of seam cell duplications in wild isolates of *C. elegans*. **(A)** Plot showing average seam cell number at 20° and 25° in N2 (depicted in green), JU775 (in purple), JU2519 (in pink), JU2007 (in light blue), XZ1516 (in yellow) and CB4856 (in orange). Error bars show 95% confidence interval of the mean. **(B)** Quantification of seam cell duplications in these isolates at 20° and 25°. Note that some isolates are more sensitive (JU2519, XZ1516) or less sensitive (CB4856) than N2 to display temperature-driven seam cell duplications. Statistical comparisons are within strains between 20 and 25° (*** *P* <0.001 with t-test (A) or binomial test (B), n≥120 for all strains).

## Discussion

### Higher temperatures lead to reproducible cell fate changes in the epidermis

Changes in temperature can have remarkable effects on both the physiology and development of organisms. For example, temperature decrease is known to extend lifespan and a recent study in *C. elegans* showed that this acts through reduction of age-mediated exhaustion of germline stem cells (Lee *et al*. 2019). Temperature can affect core cellular processes, such as the timing of cell division, which may be directly linked to temperature-imposed fitness barriers (Begasse *et al*. 2015). We report here that increasing growth temperature, yet within the physiological range of temperatures for *C. elegans* culture, leads to a gradual increase in the average number of seam cells. Interestingly, additional seam cells do not occur at random along the body axis of the animal, but seam lineages display differential sensitivity. For example, we report that V6a, V5, V1 and V2 cells are hotspots for seam cell duplications in the N2 background. Among these cells, V6a shows the highest frequency of seam cell duplications, which is in contrast to its almost insensitive posterior neighbour V6p. This is of note because V6a and V6p share the same developmental history until their common precursor divides symmetrically at the L2 stage to generate the two sub-lineages. The stereotypical distribution of seam cell duplications highlights that intrinsic differences may sensitise seam cells to respond to temperature-dependent changes affecting critical developmental signals that influence stem cell division and differentiation in the epidermis.

With regard to cell fate patterning, a well-studied example of how environmental changes can affect developmental fidelity is vulva development. Vulva formation is very robust to temperature change with the frequency of cell fate patterning errors remaining extremely low within the physiological range of temperatures, while the exact type of errors depends on the environmental pressure (Braendle and Felix 2008). For example, growth at 25° results in a low frequency (usually less than 1%) of undivided P4.p and P8.p cells, which are part of the vulval competence group although these cells do not contribute directly to the formation of the vulva by acquiring (primary or secondary) vulval cell fates (Braendle and Felix 2008). Vulva development can be more substantially de-buffered via subjecting animals to extreme non-physiological temperature challenges, such as transient growth at 30° (Grimbert and Braendle 2014). Similar to the seam, vulval cells also display cell-specific sensitivities to high temperature, with secondary vulval cells being more affected than primary cells via a decrease in Notch pathway activity (Grimbert and Braendle 2014). It will be interesting in the future to establish a mechanistic framework describing the outcome of cell fate decisions as a function of the growth temperature both within and beyond the seam cell lineage.

### Seam cell lineage defects in response to high temperature are dependent upon the Hox gene *mab-5*

The cell-specific sensitivity to duplication along the body axis at 25° led us to investigate whether Hox genes may be involved in the manifestation of this developmental phenotype. We found that the posterior Hox gene and *Antennapedia* homolog *mab-5,* plays a role in hermaphrodite seam cell patterning because it is required for posterior seam cell maintenance, as well as V6a seam cell duplications in response to temperature increase. Interestingly, expansion of the *mab-5* expression domain to the anterior end of the animal leads to a high frequency of anterior seam cell duplications even at 20°, indicating that *mab-5* may directly trigger, or at least sensitise epidermal cells to convert from any asymmetric mode of division to a symmetric one. The sensitivity in lineages such as V2, which is outside the endogenous *mab-5* expression domain but produces extra seam cells at 25°, highlights that other factors must act in parallel to *mab-5*. For example, the Hox gene *lin-39* is not required for V6a seam cell division symmetrisation at 25° but might still play a role in the less frequent anterior seam cell duplications as it mildly suppressed seam cell number increase at 25° (Figure S2A). Although *mab-5* has not been studied before in the context of hermaphrodite seam cell development, it was previously known that *mab-5* plays a role in the seam cell lineage in males, where dynamic *mab-5* expression in the V5 and V6 lineage has been shown to be required for sensory ray formation by regulating cell fate proliferation and differentiation (Salser and Kenyon 1996; Hunter *et al*. 1999).

Our findings support that *mab-5* is required for the V6a duplication errors in response to temperature presumably by creating a permissive environment within the posterior region of the animal for seam cell fate changes to occur in response to temperature increase. This is supported by the expanded range of cells displaying seam cell duplications in response to an expanded *mab-5* expression domain. Notably, *mab-5* may act in a cell-autonomous manner as mRNA expression was detected in anterior daughter cells before these differentiate to become hyp7. Interestingly, *mab-5* mRNA levels showed a maximum at V6a, which is the very same cell that also displays the highest frequency of symmetrisation at 25°. This raises the possibility that *mab-5* levels may relate to the enhanced sensitivity of V6a cells with respect to undergoing a cell fate change. Higher temperatures are thought to generally increase levels of gene expression, but this relationship varies from one gene to another (Gomez-Orte *et al*. 2018). We found no evidence to suggest that an increase in temperature triggers an increase in the expression of *mab-5.* Taken together, these results are consistent with a model in which *mab-5* is required for the symmetrisation of division at 25°, although these errors do not arise due to changes in the levels of *mab-5* transcription. It is interesting that *mab-5* has been previously reported to be involved in the competence of other epidermal cells to respond to developmental signals, namely the ventral epidermal precursor cells in males, the most posterior of which give rise to the hook sensillum group [P(9-11).p]. In this context, *mab-5* is necessary for P(9-11).p cell specification and overexpression in anterior P(1-8).p cells makes them competent to generate posterior epidermal fates depending on the activity of the Notch and EGF pathway (Yu *et al*. 2010).

### Seam cell fate changes in response to temperature increase may be due to both induced and intrinsic differences in Wnt pathway asymmetry

We present here evidence that symmetrisation of Wnt pathway activity may underlie the seam cell defects in response to growth at higher temperatures. First, we found that anterior V6a seam cell daughters show higher *egl-18* expression values at 25° than their posterior counterparts and at a frequency that matches the penetrance of the seam cell duplication phenotype, while the posterior cells exhibited overall decreased expression at 25° compared to 20°. This pattern of *egl-18* expression symmetrisation has been previously reported in *lin-22* mutants, which also show stochastic symmetrisation of seam cell divisions along the body axis at late larval stages (Katsanos *et al*. 2017). In addition to *egl-18* expression, low POP-1 levels are generally thought to correlate with higher potential of Wnt pathway activation. We found an overall decrease in nuclear POP-1 levels at 25° in all V cells, which is consistent with the higher probability of finding additional seam cells. Interestingly, V6a daughter cells appeared more symmetric in POP-1 levels compared to V6p daughters both at 20 and 25° at the L4 stage, which may explain the higher sensitivity of V6a cells to increased temperature.

With regard to upstream signalling components, we found that *bar-1* suppressed the seam cell lineage defect at 25°. This is unexpected because asymmetric seam cell divisions are thought to depend on the beta-catenins WRM-1 and SYS-1, which regulate POP-1 subcellular redistribution and transcriptional activity respectively (Rocheleau *et al*. 1999; Lo *et al*. 2004; Shetty *et al*. 2005; Phillips *et al*. 2007)The potential contribution of WRM-1 and SYS-1 to the extra seam cells observed in response to temperature was not investigated due to the temperature sensitivity of the mutants available and their strong seam cell defects. While *bar-1* is known to modulate Wnt expression through the canonical molecular pathway (Sawa and Korswagen 2013), its role in seam cell development has been questioned, partly because of the very mild seam cell defects in *bar-1* loss-of-function mutants at 20° (Figure S2B). However, we have found that *bar-1* is expressed in anterior and posterior seam cells and facilitates the seam cell symmetrisation phenotype at 25°. The activation of *bar-1* has been shown to increase Wnt target genes such as *egl-18* in the epidermis (Gorrepati *et al*. 2013; Jackson *et al*. 2014). This raises the possibility that ectopic and *bar-1* dependent activation of the Wnt pathway in the anterior V6a daughter may underlie the seam cell symmetrisation phenotype. It will be interesting to dissect in the future how increased temperature leads to this activation, with our results highlighting that intrinsic differences in Wnt pathway asymmetry may sensitise specific V cells to divide symmetrically.

### Background genetic variation influences seam cell development

Cryptic genetic variation refers to genetic variation that is phenotypically silent under wild-type conditions, yet it can have phenotypic consequences when a biological system is perturbed (Gibson and Dworkin 2004; Paaby and Rockman 2014). Cryptic genetic variation can therefore remain neutral in populations but become adaptive or deleterious upon perturbation, which is thought to relate to the increased prevalence of certain human diseases in modern times (Gibson and Dworkin 2004; Gibson and Reed 2008). Cryptic genetic variation can be revealed in model organisms via system perturbations, such as introducing mutations into divergent wild isolate backgrounds or subjecting animals to various environmental treatments. Recent efforts have therefore succeeded in detecting cryptic genetic variation affecting molecular (Snoek *et al*. 2017) or developmental processes in *C. elegans* including embryogenesis (Paaby *et al*. 2015), germ layer specification (Torres Cleuren *et al*. 2019) and vulva development (Milloz *et al*. 2008; Duveau and Felix 2012; Grimbert and Braendle 2014).

We reveal here for the first time cryptic genetic variation underlying seam cell development by demonstrating that divergent wild *C. elegans* isolates show significant differences in seam cell number when grown at 25°, while they show a comparable average seam cell number when grown at the standard growth temperature of 20°. Our results suggest that differences in the frequency of seam cell duplications over various lineages among isolates at 25° largely account for the differences in mean seam cell number. Among all isolates, XZ1516 is the most sensitive strain and displays seam cell duplications in various cell lineages at 25°. On the other hand, CB4856 is the least sensitive strain and its average seam cell number was not significantly affected by temperature. It is currently unclear if CB4856 also shows tolerance to temperature increase in other development contexts, however, this strain has been reported to show a preference for colder temperature (Anderson *et al*. 2007). Interestingly, XZ1516 and CB4856 were both sampled at the same geographic location (Hawaii), which highlights that the evolution of this developmental phenotype is unlikely to reflect some specific ecological adaptation. It is also of note that certain isolates displayed changes in the frequencies of seam cell duplication both within or outside the range of cells that are affected in the N2 background. For example, isolates JU775 and JU2007 showed enhanced sensitivity in V1a and V6a, which is similar to N2, whereas JU2519 and XZ1516 showed novel duplications in H1, H2, V3 and T cell lineages. The broad expansion of cell-specific sensitivity observed in strain XZ1516 is reminiscent of the *mab-5* gain of function phenotype in N2 and may also reflect evolution in key morphogenetic factors.

It will be interesting in the future to discover the genetic architecture of cryptic genetic variation underlying seam cell development. Given the importance of the Wnt signalling pathway in regulating seam cell development, it is intriguing to speculate that differences in Wnt pathway activity or sensitivity of the response to Wnt activation among isolates may underlie the cryptic genetic variation observed. Previous studies have revealed cryptic genetic variation in cell-specific Ras and Notch pathway activity outputs in the context of the vulval signalling network (Milloz *et al*. 2008). More recently, cryptic variation was detected in the contribution of the Wnt input in the gene regulatory network underlying endoderm specification (Torres Cleuren *et al*. 2019). Quantifying changes in the abundance of Wnt signalling components among isolates is likely to be challenging at the whole organism level (Singh *et al*. 2016). Seam cell development may thus offer a suitable tissue-specific read-out to facilitate the discovery of genetic modifiers influencing the conserved Wnt signalling pathway, with possible implications in human development and disease.

---

**Figure S1:**
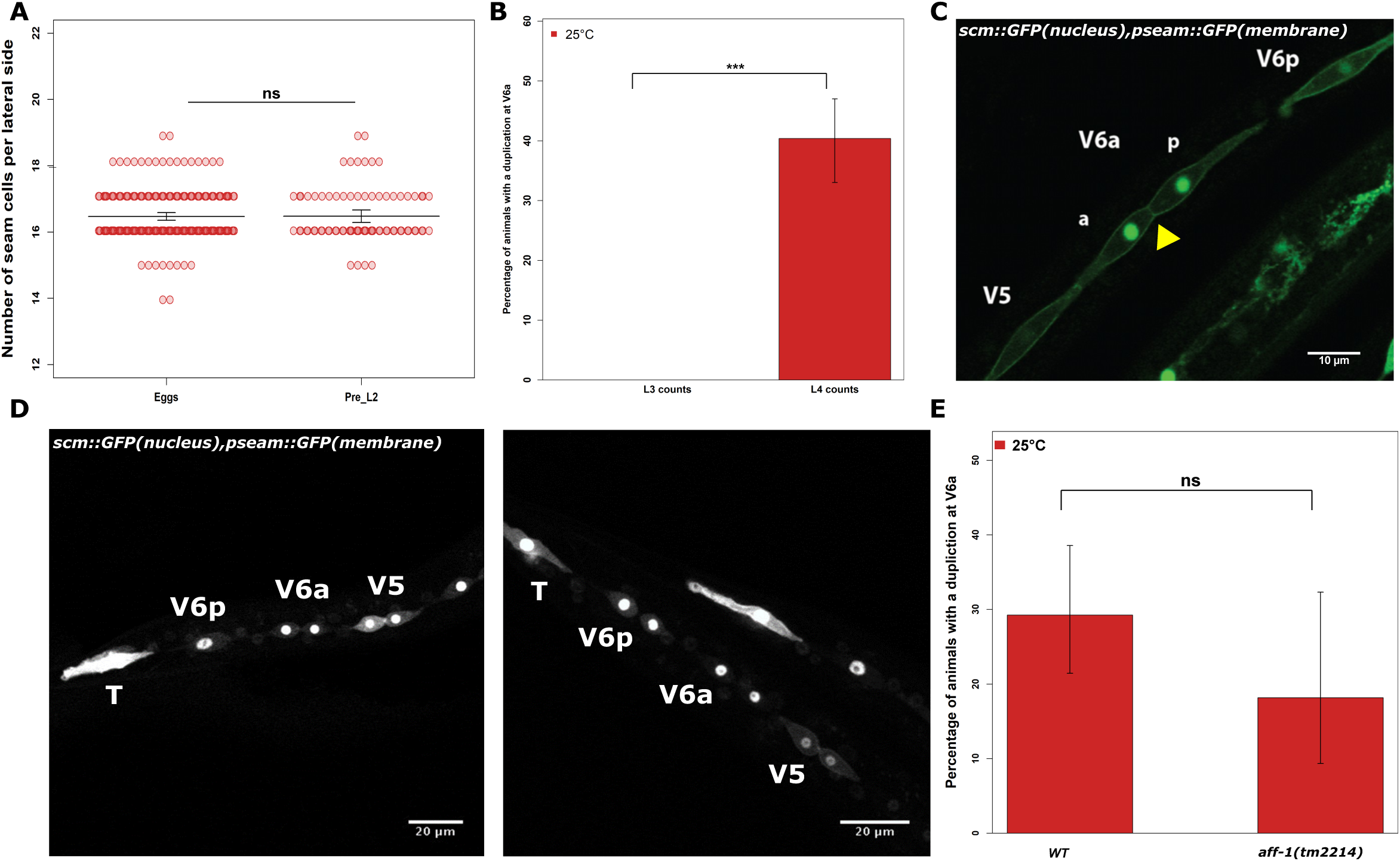
Phenotypic analysis of seam cell duplications in response to temperature increase. **(A)** Counts of seam cells from N2 (WT) animals shifted to 25° prior to egg hatching or prior to L2. In both cases, an equivalent increase in seam cell number was observed. **(B)** The frequency of V6 duplications counted at the end of the L3 and L4 larval stage (*** corresponds to *P* value <0.001, with a binomial test). **(C-D)** Representative images of posterior seam cells in s*cm::GFP; pseam::GFP::CAAX* animals, which display GFP in the nucleus and membrane of seam cells. (C) V6a duplications are not due to cytokinesis errors as cells separate normally post division. Yellow arrowhead indicates membrane boundary between V6aa and V6ap. (D) V6a is not the last cell to divide. Left image shows V6p yet to divide, while V6a and V5 are dividing, middle panel displays both V6p and V5 dividing after V6a. **(E)** V6a duplications still occur in animals lacking *aff-1* activity. Error bars indicate standard error of the proportion. Scale bars are 10 µm in C and 20 µm in D.

**Figure S2:**
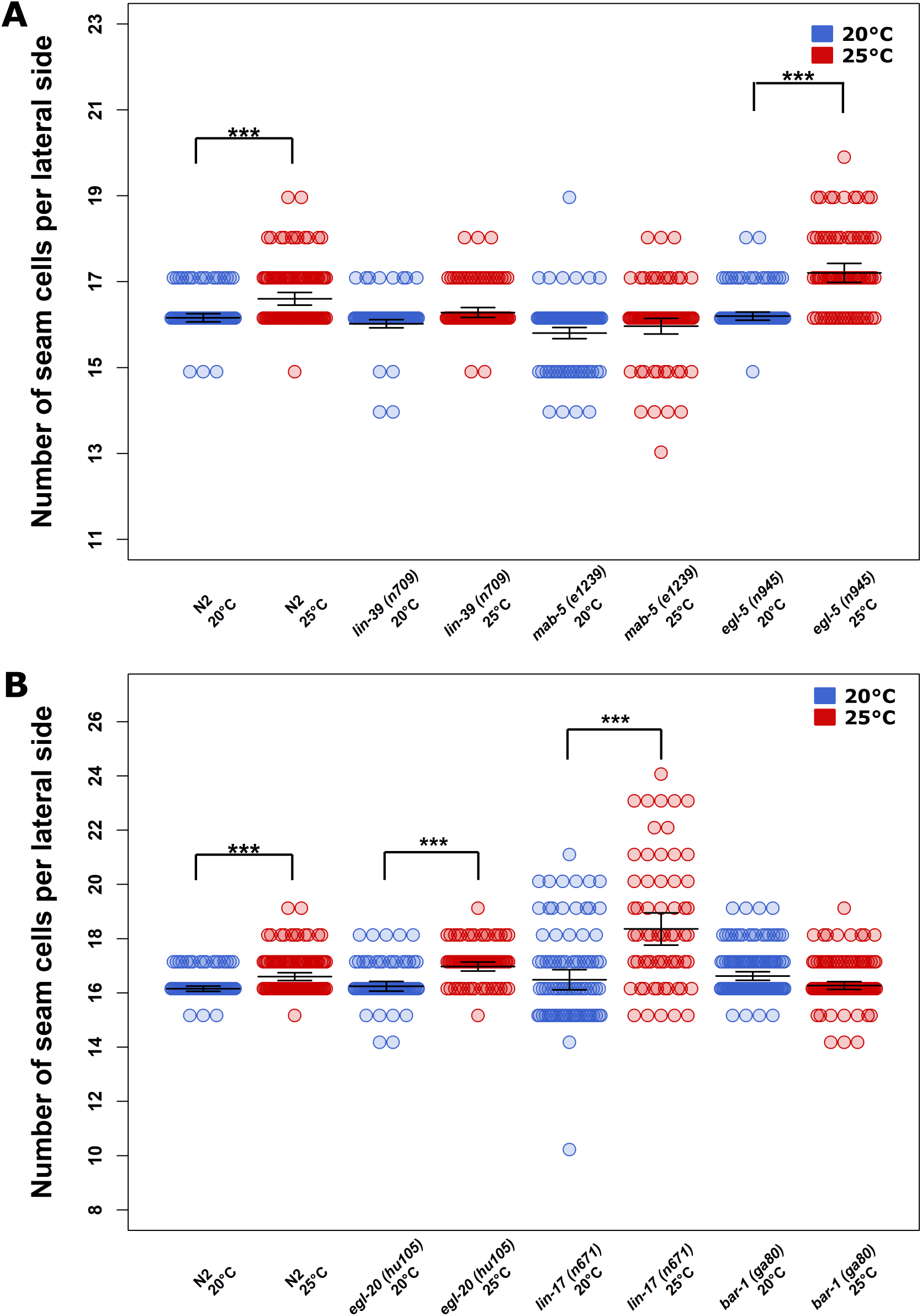
Complete seam cell counts in mutant backgrounds at 20° and 25°. **(A)** Counts of seam cells in strains deficient for the Hox genes, *lin-39(n709)*, mab-5(e1239) and *egl-5(n945)*. **(B)** Seam cell counts in strains deficient for the Wnt genes, *egl-20(hu105)*, *lin-17(n671)* and *bar-1(ga80)*. *P*, ***<0.001 with a two-sample t-test. Error bars indicate 95% confidence intervals of the mean.

**Figure S3:**
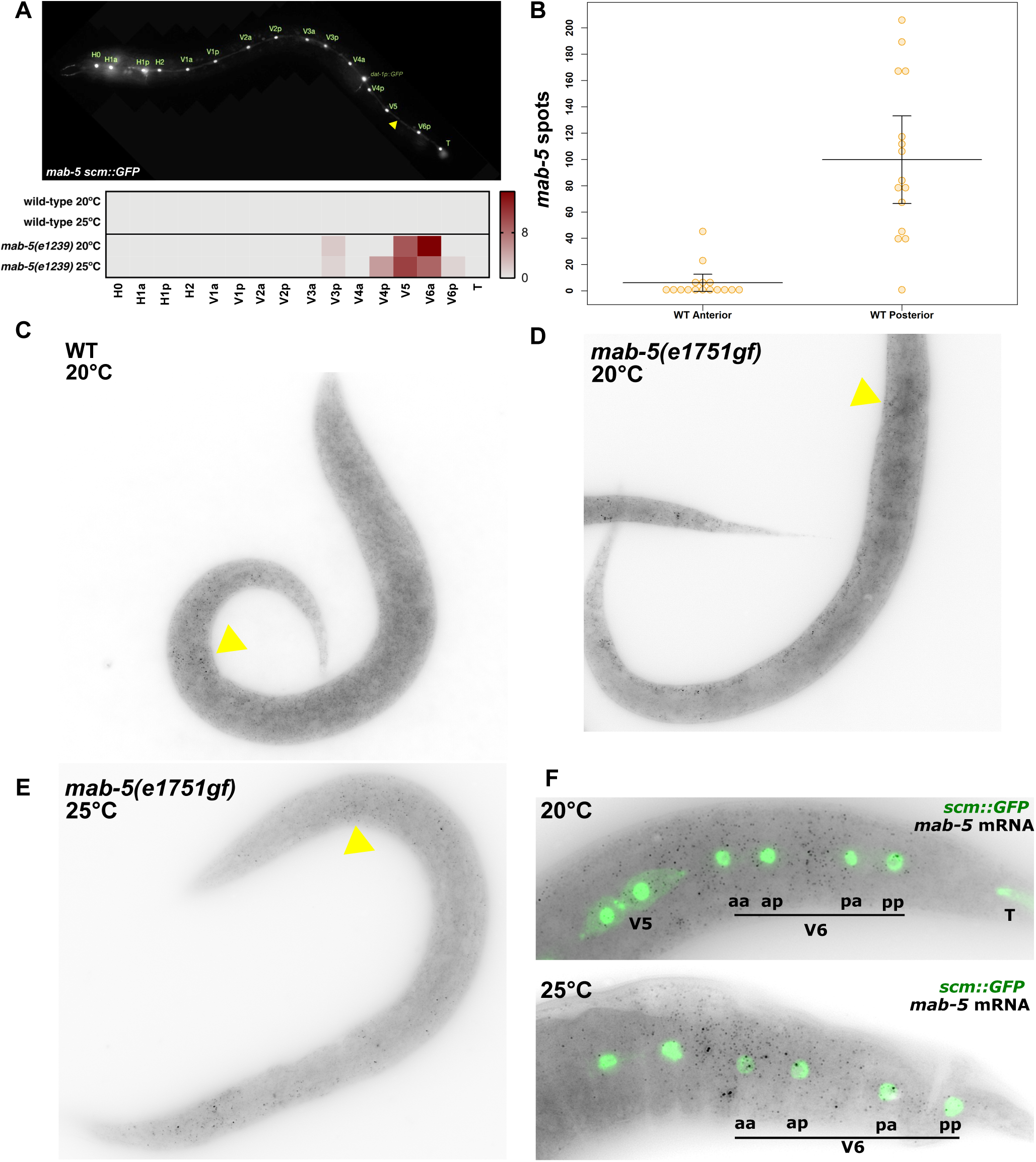
Analysis of *mab-5* expression and function in seam cells. **(A)** Posterior seam cell loss in *mab-5(e1239)* animals. Upper panel is a representative image of V6a loss (yellow arrowhead). Lower panel shows heatmap displaying frequency of seam cell loss in *mab-5(e1239)* versus WT animals at 20° and 25°. The difference in the frequency of seam cell loss between the two temperatures is not significant with a binomial test. **(B)** Counts of *mab-5* mRNA in L1 animals by smFISH (n=15). Note higher *mab-5* expression in the posterior side of the animal compared to the anterior. Anterior and posterior were defined here as before and after V3a respectively. **(C-E)** Representative images of *mab-5* smFISH in WT N2 at 20°C, *mab-5(e1751gf)* at 20° and *mab-5(e1751gf)* at 25°. Yellow arrows show ectopic anterior expression of *mab-5* in the gain of function mutant background. **(F)** Representative close-up images of *mab-5* mRNA expression at 20° and 25° in V6 cells following the L4 division.

**Figure S4:**
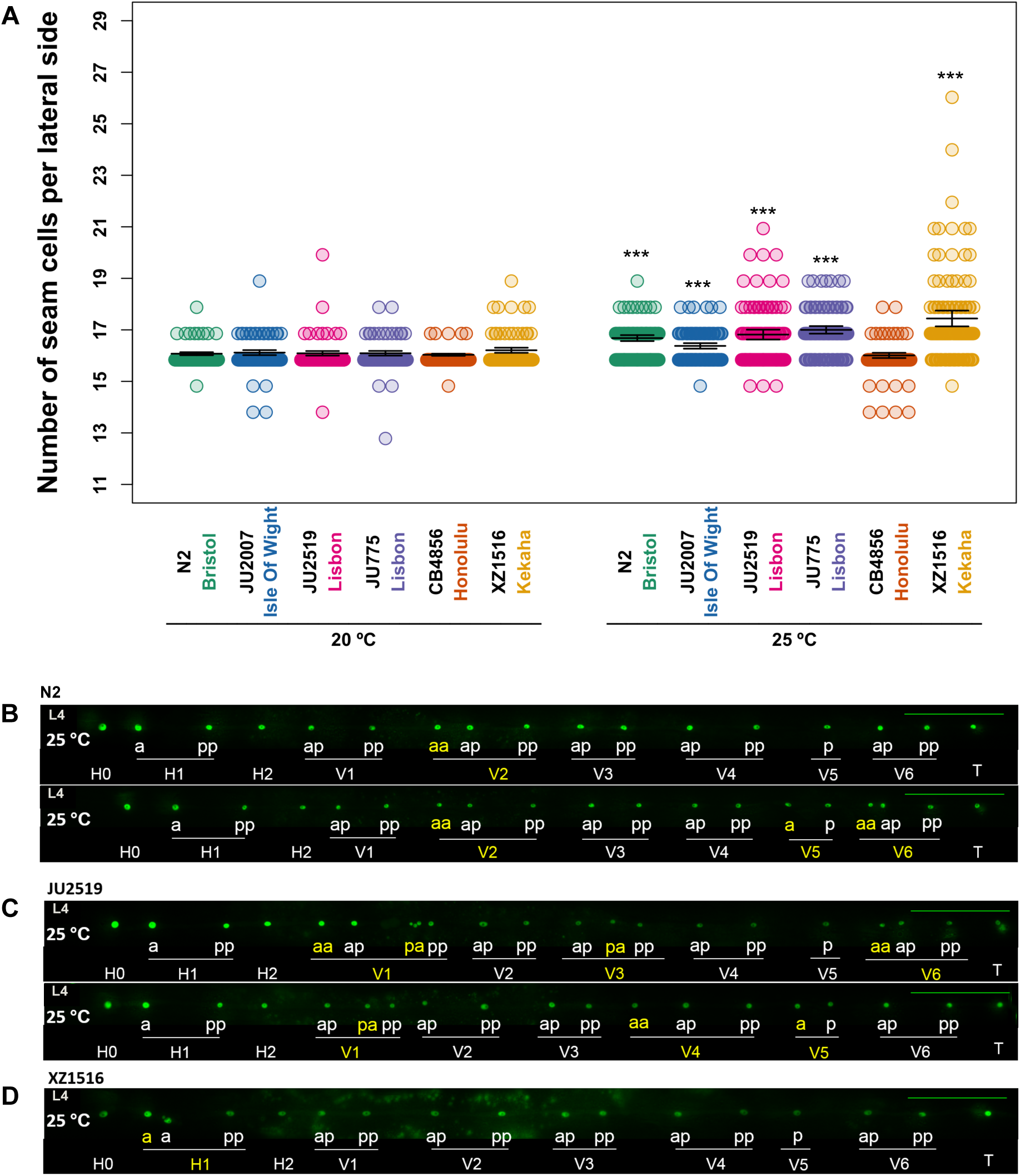
Wild isolates of *C. elegans* show differences in frequency of seam cell duplications in response to temperature increase. **(A)** Quantification of seam cells in N2 (Bristol), JU775 (Lisbon), JU2519 (Lisbon), JU2007 (Isle of Wight), XZ1516 (Kehaka) and CB4856 (Honolulu) at 20° and 25°, *** corresponds to *P*< 0.001, two-sample t-test comparisons between same isolate at 20° and 25°. Error bars indicate 95% confidence intervals of the mean. **(B**) Representative images of errors in N2 animals at L4 grown at 20° and 25°, yellow labels indicate extra seam cells in V2, V5 and V6. **(C)** Representative images of errors in JU2519 grown at 25°, yellow labels indicate extra seam cells in V1, V3, V4, V5 and V6. **(D)** Representative images of errors in XZ1516 grown at 25°, yellow labels indicate extra seam cells in H1. Scale bar in green is 100 µm.

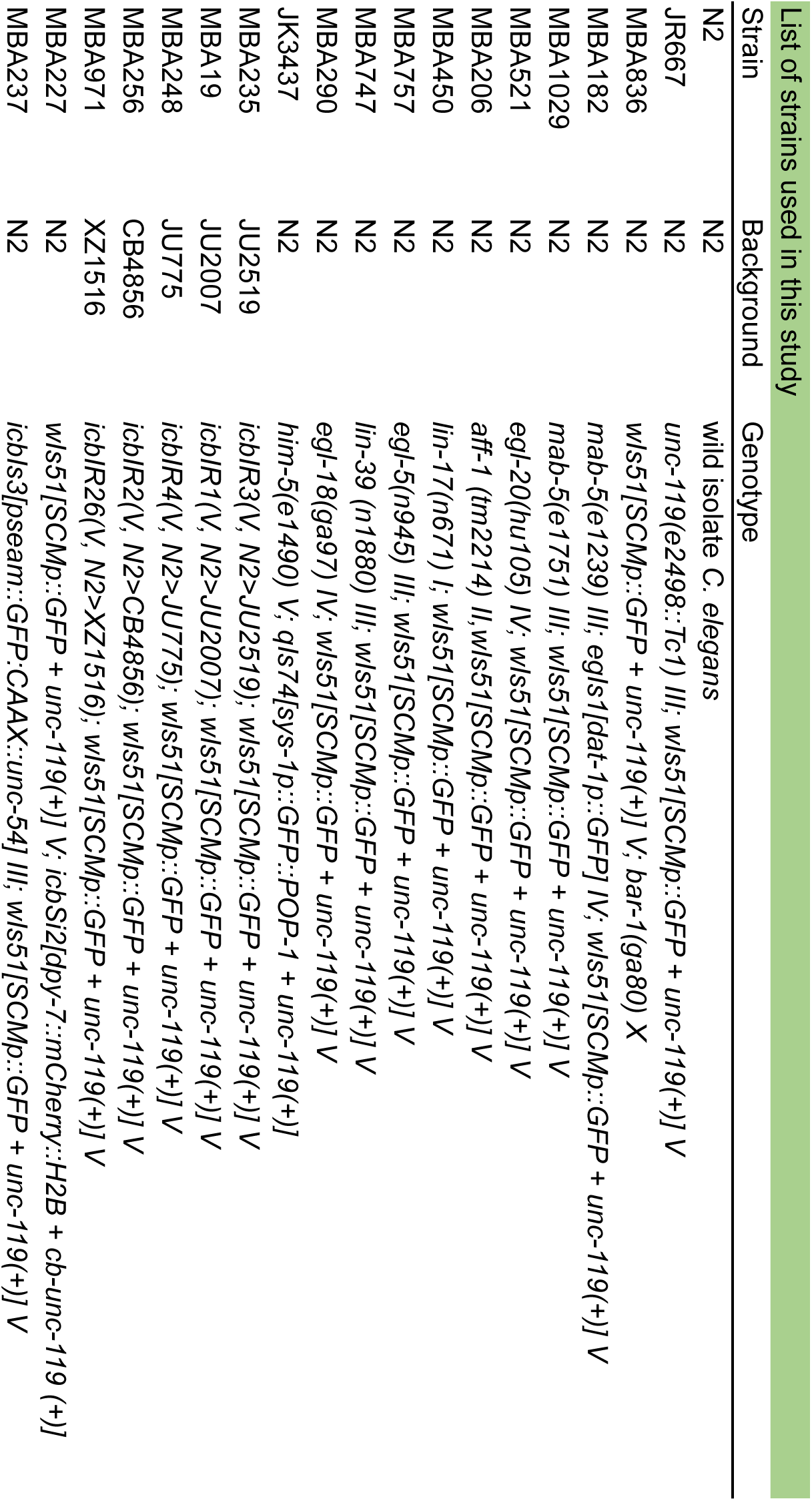

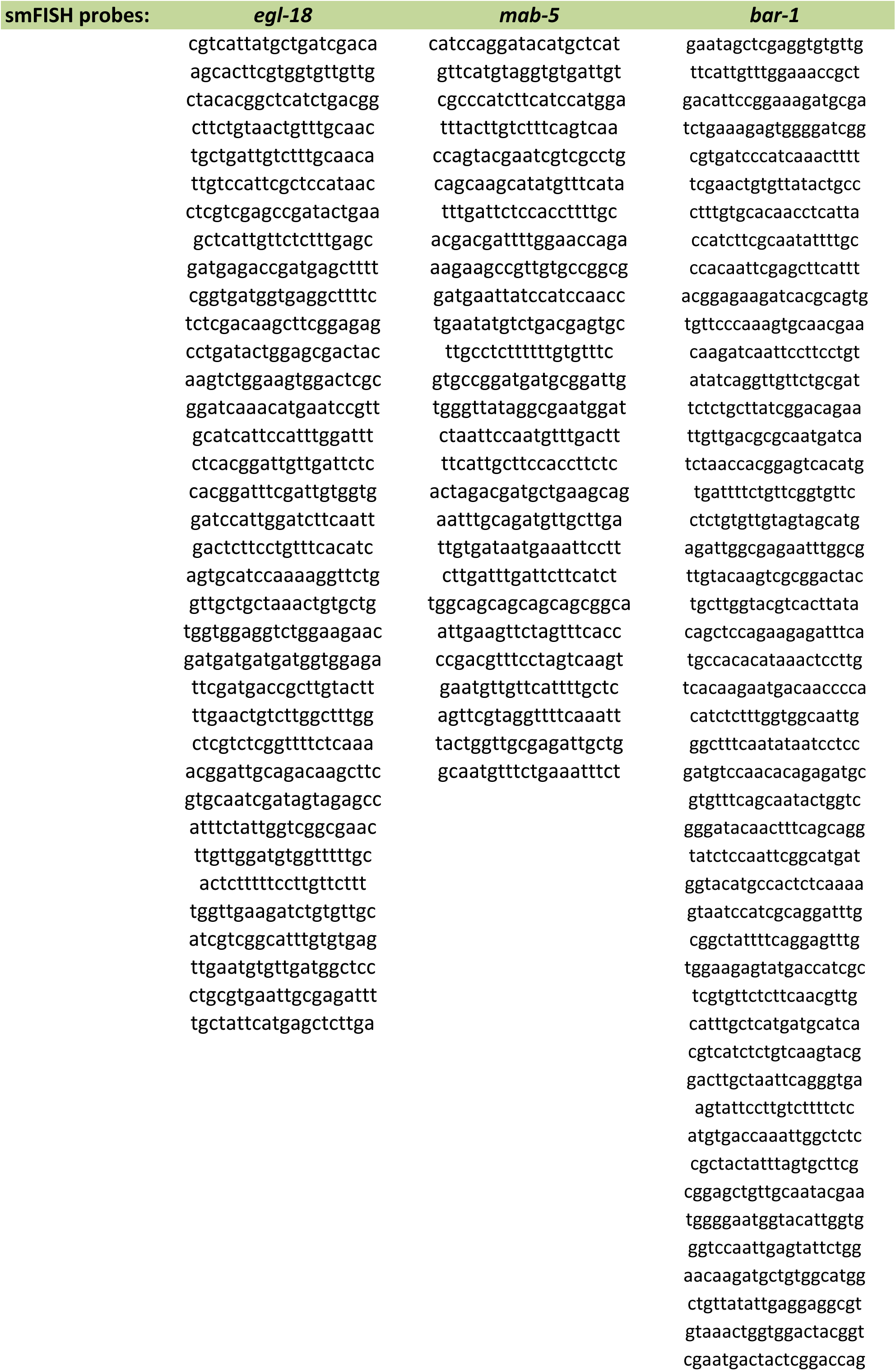

*mab-5* probe
| 20°C |  |  |  |  |  | 25°C |  |  |  |  |  |
| --- | --- | --- | --- | --- | --- | --- | --- | --- | --- | --- | --- |
| V5a | V5p | V6aa | V6ap | V6pa | V6pp | V5a | V5p | V6aa | V6ap | V6pa | V6pp |
| 20 | 6 | 19 | 3 | 11 | 2 | 7 | 9 | 16 | 14 | 0 | 5 |
| 7 | 1 | 46 | 8 | 10 | 20 | 4 | 4 | 21 | 7 | 32 | 17 |
| 4 | 24 | 25 | 25 | 25 | 9 | 4 | 13 | 4 | 4 | 13 | 9 |
| 0 | 1 | 18 | 12 | 30 | 7 | 5 | 2 | 34 | 5 | 17 | 6 |
| 14 | 11 | 19 | 7 | 2 | 5 | 4 | 2 | 9 | 4 | 8 | 29 |
| 0 | 6 | 23 | 5 | 37 | 19 | 0 | 20 | 17 | 10 | 15 | 1 |
| 2 | 8 | 26 | 28 | 12 | 13 | 0 | 9 | 25 | 14 | 4 | 16 |
| 6 | 3 | 42 | 6 | 14 | 24 | 0 | 2 | 25 | 10 | 10 | 6 |
| 0 | 8 | 17 | 4 | 33 | 1 | 3 | 2 | 23 | 0 | 11 | 9 |
| 6 | 18 | 19 | 21 | 15 | 7 | 12 | 30 | 22 | 8 | 3 | 29 |
| 1 | 14 | 44 | 4 | 17 | 7 | 6 | 13 | 29 | 17 | 30 | 7 |
| 6 | 6 | 40 | 18 | 11 | 5 | 5 | 3 | 18 | 10 | 18 | 18 |
| 2 | 5 | 20 | 27 | 22 | 0 | 0 | 36 | 9 | 1 | 11 | 2 |
| 4 | 3 | 31 | 8 | 10 | 0 | 9 | 22 | 14 | 3 | 0 | 15 |
| 7 | 8 | 54 | 14 | 50 | 10 | 0 | 29 | 14 | 18 | 6 | 8 |
| 1 | 13 | 13 | 28 | 16 | 17 | 1 | 10 | 29 | 16 | 3 | 28 |
| 5 | 5 | 9 | 19 | 13 | 5 | 8 | 10 | 16 | 6 | 8 | 34 |
| 6 | 14 | 45 | 24 | 16 | 0 | 1 | 1 | 3 | 0 | 0 | 40 |
| 9 | 11 | 21 | 10 | 4 | 12 | 2 | 4 | 32 | 17 | 11 | 17 |
| 3 | 2 | 20 | 5 | 19 | 3 | 2 | 16 | 5 | 5 | 2 | 15 |
| 20 | 28 | 57 | 27 | 40 | 0 | 12 | 25 | 15 | 2 | 0 | 28 |
| 11 | 2 | 39 | 25 | 10 | 28 | 10 | 23 | 25 | 7 | 18 | 20 |
| 12 | 31 | 18 | 27 | 3 | 5 | 0 | 6 | 24 | 8 | 9 | 23 |
| 3 | 2 | 41 | 19 | 19 | 24 | 8 | 0 | 22 | 10 | 14 | 3 |
| 21 | 0 | 45 | 21 | 29 | 13 | 3 | 4 | 28 | 28 | 23 | 1 |
| 14 | 4 | 25 | 26 | 30 | 14 | 17 | 7 | 25 | 11 | 4 | 8 |
| 2 | 17 | 11 | 3 | 11 | 42 | 1 | 12 | 20 | 12 | 36 | 15 |
| 9 | 7 | 26 | 15 | 7 | 16 | 4 | 15 | 39 | 21 | 29 | 3 |
| 0 | 9 | 16 | 17 | 3 |  |  |  |  |  |  | 11 |

**bar-1** probe
| 20°C |  |  |  | 25°C |  |  |  |
| --- | --- | --- | --- | --- | --- | --- | --- |
| V6aa | V6ap | V6pa | V6pp | V6aa | V6ap | V6pa | V6pp |
| 11 | 3 | 13 | 1 | 24 | 6 | 4 | 18 |
| 6 | 9 | 18 | 7 | 12 | 13 | 18 | 14 |
| 11 | 5 | 9 | 5 | 6 | 17 | 4 | 8 |
| 4 | 5 | 2 | 12 | 15 | 16 | 4 | 6 |
| 10 | 1 | 12 | 4 | 19 | 3 | 4 | 1 |
| 3 | 13 | 2 | 5 | 17 | 10 | 9 | 16 |
| 12 | 18 | 9 | 4 | 10 | 6 | 0 | 2 |
| 14 | 2 | 9 | 0 | 5 | 9 | 3 | 9 |
| 14 | 9 | 10 | 12 | 2 | 6 | 14 | 7 |
| 2 | 11 | 3 | 9 | 12 | 9 | 6 | 6 |
| 8 | 16 | 8 | 8 | 2 | 8 | 1 | 1 |
| 1 | 4 | 11 | 5 | 5 | 10 | 9 | 3 |
| 10 | 0 | 6 | 4 | 20 | 3 | 20 | 2 |
| 1 | 3 | 9 | 2 | 0 | 11 | 1 | 23 |
| 7 | 5 | 10 | 9 | 0 | 6 | 3 | 2 |
| 5 | 0 | 5 | 1 | 2 | 2 | 5 | 1 |
| 8 | 8 | 9 | 7 | 12 | 3 | 7 | 1 |
| 3 | 8 | 1 | 11 | 0 | 6 | 1 | 7 |
| 26 | 1 | 29 | 4 | 3 | 0 | 2 | 1 |
| 7 | 3 | 12 | 4 | 7 | 1 | 7 | 1 |
| 7 | 7 | 5 | 23 | 22 | 16 | 13 | 4 |
| 9 | 10 | 9 | 6 | 15 | 33 | 12 | 12 |
| 2 | 2 | 4 | 3 | 9 | 26 | 14 | 12 |
| 6 | 16 | 9 | 6 | 17 | 26 | 12 | 14 |
| 9 | 3 | 12 | 2 | 35 | 36 | 8 | 12 |
| 4 | 3 | 13 | 9 | 5 | 35 | 4 | 11 |
| 7 | 7 | 16 | 6 | 7 | 29 | 16 | 3 |
| 23 | 7 | 22 | 14 | 4 | 7 | 9 | 5 |
|  | 10 |  | 9 | 12 | 5 | 6 | 7 |
|  | 19 |  | 12 | 1 | 9 | 8 | 2 |
|  |  |  |  | 4 | 4 | 4 | 4 |
|  |  |  |  | 12 | 1 | 6 | 5 |
|  |  |  |  | 5 | 1 | 3 | 3 |
|  |  |  |  | 6 | 9 | 24 | 7 |
|  |  |  |  | 2 | 7 | 1 | 14 |
|  |  |  |  | 9 | 2 | 4 | 1 |
|  |  |  |  | 3 | 23 | 3 | 4 |
|  |  |  |  | 5 | 1 | 17 | 7 |
|  |  |  |  | 0 | 8 | 22 | 8 |
|  |  |  |  | 6 | 0 | 5 | 6 |
|  |  |  |  | 16 | 4 | 11 | 5 |
|  |  |  |  | 15 | 13 | 3 | 3 |
|  |  |  |  | 6 | 11 | 1 | 2 |
|  |  |  |  | 9 | 8 | 11 | 6 |
|  |  |  |  | 14 | 9 | 1 | 15 |
|  |  |  |  | 1 | 10 | 11 | 0 |
|  |  |  |  | 2 | 1 | 2 | 9 |
|  |  |  |  | 13 | 0 | 1 | 3 |
|  |  |  |  | 8 | 5 | 0 | 3 |
|  |  |  |  | 2 | 10 | 7 | 0 |
|  |  |  |  | 0 | 0 | 1 | 5 |
|  |  |  |  | 2 | 0 |  | 1 |
|  |  |  |  | 8 | 2 |  |  |
|  |  |  |  |  | 7 |  |  |

*egl-18* probe
| 20°C |  |  |  |  |  | 25°C |  |  |  |  |  |
| --- | --- | --- | --- | --- | --- | --- | --- | --- | --- | --- | --- |
| V5a | V5p | V6aa | V6ap | V6pa | V6pp | V5a | V5p | V6aa | V6ap | V6pa | V6pp |
| 10 | 54 | 21 | 54 | 12 | 4 | 8 | 56 | 5 | 7 | 23 | 26 |
| 4 | 41 | 6 | 39 | 11 | 27 | 16 | 14 | 1 | 3 | 16 | 11 |
| 7 | 43 | 4 | 80 | 8 | 74 | 44 | 26 | 43 | 32 | 6 | 41 |
| 1 | 59 | 5 | 69 | 9 | 59 | 8 | 47 | 8 | 39 | 0 | 1 |
| 4 | 35 | 3 | 34 | 16 | 25 | 7 | 37 | 57 | 54 | 2 | 23 |
| 5 | 47 | 13 | 54 | 9 | 47 | 1 | 12 | 12 | 19 | 0 | 21 |
| 4 | 33 | 0 | 21 | 3 | 6 | 4 | 65 | 2 | 49 | 2 | 17 |
| 14 | 62 | 34 | 49 | 11 | 58 | 9 | 31 | 2 | 40 | 0 | 6 |
| 0 | 22 | 5 | 4 | 6 | 58 | 7 | 28 | 37 | 32 | 4 | 47 |
| 1 | 44 | 3 | 38 | 5 | 45 | 0 | 14 | 4 | 15 | 2 | 7 |
| 4 | 33 | 9 | 13 | 2 | 26 | 11 | 39 | 2 | 20 | 1 | 30 |
| 2 | 33 | 4 | 47 | 1 | 54 | 0 | 34 | 3 | 22 | 0 | 50 |
| 0 | 61 | 2 | 58 | 0 | 43 | 6 | 15 | 3 | 6 | 2 | 8 |
| 11 | 74 | 9 | 77 | 17 | 50 | 1 | 13 | 3 | 22 | 3 | 27 |
| 0 | 62 | 2 | 15 | 0 | 23 | 7 | 16 | 6 | 31 | 10 | 15 |
| 0 | 28 | 1 | 33 | 11 | 29 | 0 | 28 | 9 | 28 | 4 | 19 |
| 2 | 12 | 8 | 19 | 1 | 29 | 4 | 16 | 0 | 6 | 3 | 27 |
| 2 | 12 | 2 | 33 | 4 | 34 | 0 | 11 | 0 | 22 | 1 | 29 |
| 18 | 21 | 13 | 49 | 6 | 32 | 3 | 4 | 0 | 3 | 3 | 5 |
| 9 | 41 | 6 | 47 | 2 | 44 |  |  | 2 | 18 | 4 | 5 |
| 2 | 25 | 15 | 48 | 1 | 29 |  |  |  |  |  |  |
| 12 | 49 | 13 | 49 | 17 | 28 |  |  |  |  |  |  |
| 1 | 58 | 1 | 39 | 15 | 52 |  |  |  |  |  |  |
| 2 | 37 | 6 | 30 | 3 | 56 |  |  |  |  |  |  |
| 5 | 34 | 2 | 10 | 0 | 19 |  |  |  |  |  |  |
| 2 | 28 | 2 | 29 | 1 | 31 |  |  |  |  |  |  |
| 3 | 33 | 6 | 56 | 0 | 49 |  |  |  |  |  |  |
| 0 | 7 | 7 | 43 | 3 | 52 |  |  |  |  |  |  |
| 4 | 46 | 6 | 37 | 3 | 44 |  |  |  |  |  |  |
| 1 | 43 | 0 | 32 | 2 | 27 |  |  |  |  |  |  |
| 4 | 36 | 7 | 40 | 6 | 28 |  |  |  |  |  |  |
| 5 | 52 | 10 | 36 | 1 | 21 |  |  |  |  |  |  |

***mab-5*** probe 20°C
| anterior | posterior |
| --- | --- |
| 5 | 167 |
| 0 | 45 |
| 47 | 113 |
| 1 | 40 |
| 0 | 187 |
| 0 | 117 |
| 3 | 67 |
| 1 | 0 |
| 0 | 78 |
| 2 | 41 |
| 0 | 82 |
| 9 | 208 |
| 21 | 166 |
| 1 | 104 |
| 8 | 81 |
| 0 |  |

## Acknowledgements

We thank Julien Lambert, Michael Fasseas and Iqrah Razzaq for help with experiments. Some strains were provided by the CGC, which is funded by NIH Office of Research Infrastructure Programs (P40 OD010440). This work was supported by the European Research Council (639485-ROBUSTNET) and the Leverhulme Trust (RPG-2015-235).

